# Impacts of Taxon-Sampling Schemes on Bayesian Molecular Dating under the Unresolved Fossilized Birth-Death Process

**DOI:** 10.1101/2021.11.16.468757

**Authors:** Arong Luo, Chi Zhang, Qing-Song Zhou, Simon Y.W. Ho, Chao-Dong Zhu

**Affiliations:** Key Laboratory of Zoological Systematics and Evolution, Institute of Zoology, Chinese Academy of Sciences, Beijing 100101, China; Key Laboratory of Vertebrate Evolution and Human Origins, Institute of Vertebrate Paleontology and Paleoanthropology, Chinese Academy of Sciences, Beijing 100044, China; Center for Excellence in Life and Paleoenvironment, Chinese Academy of Sciences, Beijing 100044, China; School of Life and Environmental Sciences, University of Sydney, Sydney, New South Wales 2006, Australia; State Key Laboratory of Integrated Pest Management, Institute of Zoology, Chinese Academy of Sciences, Beijing 100101, China; College of Life Sciences, University of Chinese Academy of Sciences, Beijing, 100049, China; International College, University of Chinese Academy of Sciences, Beijing, 100049, China

**Keywords:** tip dating, total-evidence dating, fossilized birth-death process, taxon sampling, phylogenomics, eutherian mammals

## Abstract

Evolutionary timescales can be estimated using a combination of genetic data and fossil evidence based on the molecular clock. Bayesian phylogenetic methods such as tip dating and total-evidence dating provide a powerful framework for inferring evolutionary timescales, but the most widely used priors for tree topologies and node times often assume that present-day taxa have been sampled randomly or exhaustively. In practice, taxon sampling is often carried out so as to include representatives of major lineages, such as orders or families. We examined the impacts of these diversified sampling schemes on Bayesian molecular dating under the unresolved fossilized birth-death (FBD) process, in which fossil taxa are topologically constrained but their exact placements are not inferred. We used synthetic data generated by simulation of nucleotide sequence evolution, fossil occurrences, and diversified taxon sampling. Our analyses show that increasing sampling density does not substantially improve divergence-time estimates under benign conditions. However, when the tree topologies were fixed to those used for simulation or when evolutionary rates varied among lineages, the performance of Bayesian tip dating improves with sampling density. By exploring three situations of model mismatches, we find that including all relevant fossils without pruning off those inappropriate for the FBD process can lead to underestimation of divergence times. Our reanalysis of a eutherian mammal data set confirms some of the findings from our simulation study, and reveals the complexity of diversified taxon sampling in phylogenomic data sets. In highlighting the interplay of taxon-sampling density and other factors, the results of our study have useful implications for Bayesian molecular dating in the era of phylogenomics.

## Introduction

Estimating the timescale of the Tree of Life has been a long-standing goal in evolutionary biology. This endeavor is dominated by analyses based on the molecular clock, which began as a simple, idealized model of evolutionary rate constancy (Zuckerkandl and Pauling 1965) but has undergone considerable developments over the past six decades (Ho 2020). Nevertheless, all molecular clocks need to be calibrated using external time information, which is often obtained from fossil evidence. In this context, node-dating approaches use fossil data indirectly to inform constraints or prior densities on node ages (e.g., Drummond et al. 2006; Yang and Rannala 2006; Nguyen and Ho 2020), whereas tip-dating approaches enable fossils to be included as sampled tips in the tree (Pyron 2011; Ronquist et al. 2012).

Bayesian methods of molecular-clock dating hold particular appeal because they can integrate diverse sources of information about lineage diversification, nucleotide substitution, and other aspects of the evolutionary process into a unified framework (dos Reis et al. 2016; Bromham et al. 2018). In particular, Bayesian tip dating is able to combine molecular sequences and morphological characters in a coherent framework (i.e., total-evidence tip dating), thereby making full use of fossil evidence and eliminating the need to specify calibration prior densities (Ronquist et al. 2012; Zhang et al. 2016). It is often used in conjunction with a tree prior based on the fossilized birth-death (FBD) process (also called the resolved FBD process; O’Reilly and Donoghue 2020), which provides a model of lineage diversification that accounts for speciation, extinction, fossilization, and taxon sampling. (Stadler 2010; Zhang et al. 2016). Evaluations of tip-dating methods have demonstrated their ability to estimate evolutionary timescales with accuracy and precision (Gavryushkina et al. 2014; Zhang et al. 2016; Luo et al. 2020).

The resolved FBD process often poorly resolves the phylogenetic positions of fossil taxa, even when the relationships among extant taxa are reconstructed with confidence (Zhang et al. 2016; Luo et al. 2020). This problem can perhaps be attributed to the use of poorly fitting models of morphological evolution (e.g., the typical Mk model; Lewis 2001; Goloboff et al. 2019; Gavryushkina and Zhang 2020), uncertainty in fossil ages, and/or low informativeness of morphological characters (O’Reilly et al. 2015; Luo et al. 2020; Spasojevic et al. 2021). Moreover, the extent of among-lineage rate heterogeneity and convergent evolution in morphological characters, as well as the typically high proportion of missing characters for many fossil specimens (Sansom and Wills 2013; Lee and Palci 2015; O’Reilly and Donoghue 2021), have negative impacts on estimates of evolutionary parameters. These problems could potentially contribute to overestimation of divergence times in practice (e.g., Wood et al. 2013; O’Reilly et al. 2015; Ronquist et al. 2016; Spasojevic et al. 2021).

Fortunately, Bayesian tip dating can be performed on molecular sequences alone (i.e., without morphological character data), under the unresolved FBD process. In this approach, the placement of each fossil is numerically integrated over a chosen portion of the phylogeny, and the fossil ages provide calibrating information (Gavryushkina et al. 2014; Heath et al. 2014; Barido-Sottani et al. 2019). The unresolved FBD process allows a larger amount of fossil data to be used, while removing the need for extensive morphological character data from the taxa of interest. It can also address some persistent issues such as uncertainty in fossil ages (Barido-Sottani et al. 2019) and the incompleteness of fossil sampling (O’Reilly and Donoghue 2020).

The FBD model typically embeds a continuous process of fossil preservation and sampling through time (modeled by a Poisson process) in the lineage diversification history (modeled by a birth-death process of speciation and extinction) (Stadler 2010; Heath et al. 2014). Accordingly, the unresolved FBD process can account for a lineage diversification history where all of the extant taxa in the group have been completely sampled. In reality, however, sampling of extant taxa is rarely exhaustive but is often designed to include representatives of certain taxonomical ranks such as families and orders (Matschiner 2019). This diversified sampling strategy has been adopted by a range of genome-sequencing initiatives (e.g., Jarvis et al. 2014; Lewin et al. 2018; Fan et al. 2020). For example, the Bird 10,000 Genomes project (Jarvis et al. 2014) has taken a four-phase approach that involves sampling major branches (orders and families) before proceeding to the twigs (genera and species). Zhang et al. (2016) extended the FBD process to account for such intentionally diversified sampling of extant taxa, whereby exactly one extant descendant is sampled for each branch present at a certain time slice (Fig. 1; Höhna et al. 2011). This approach thus provides an opportunity to examine the potential impacts of sampling at different phylogenetic scales on inferences of macroevolutionary parameters, how taxon-sampling density interacts with other factors (e.g., choice of priors), and how fossil information should be combined with genomic data in molecular dating.

**Figure 1.**
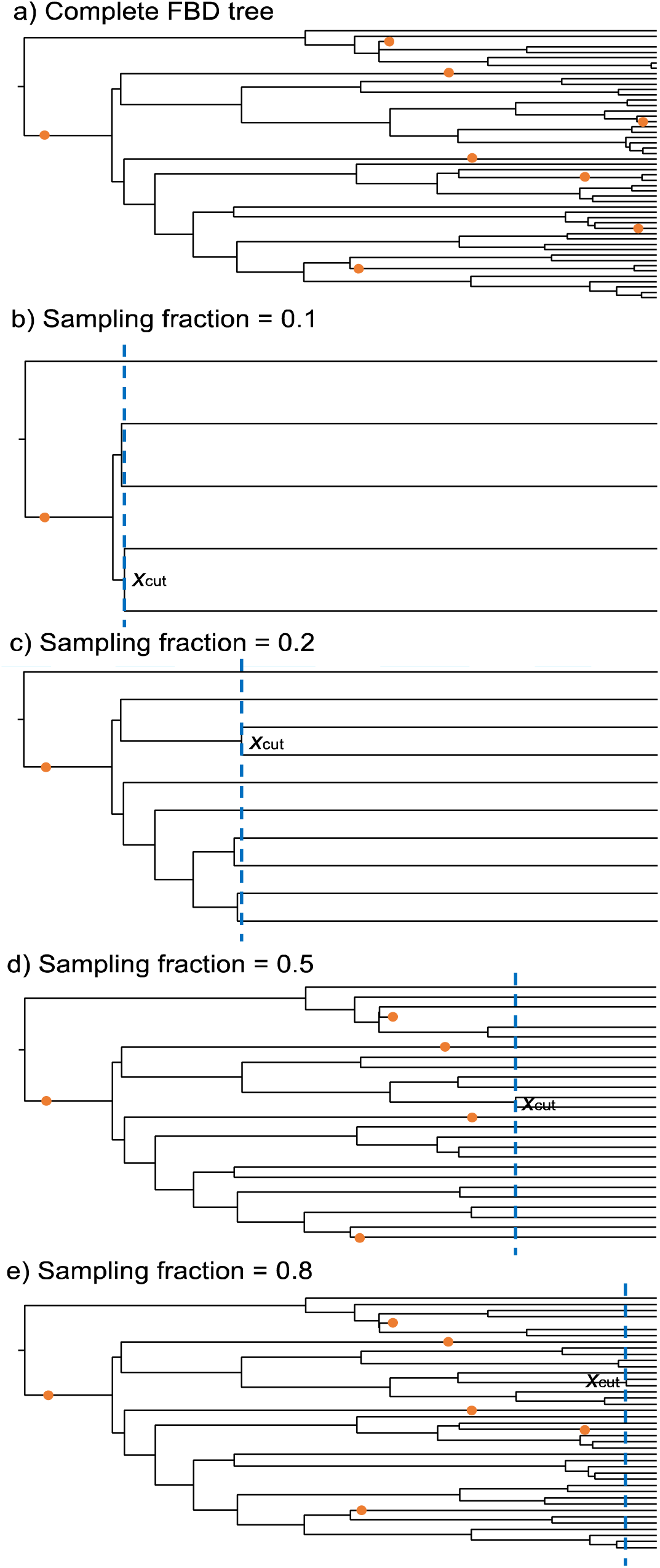
(a) A fossilized birth-death (FBD) tree depicting the reconstructed history of all present-day species (the number of extant taxa *N* = 50) and all sampled fossil occurrences (denoted by solid circles). Its original species tree was generated under the birth-death process controlled by speciation rate λ, extinction rate μ, and time of the most recent common ancestor of all extant and extinct species *t*_mrca_, based on which fossil occurrences and extant taxa were sampled by fossil recovery rate ø and sampling fraction ρ (ρ = 1 for the complete FBD tree), respectively. (b) Resultant FBD tree after diversified sampling with ρ = 0.1 and pruning fossils younger than the cut-off time *x*_cut_ (dashed vertical line). The *x*_cut_ is equal to the (*n* – 1)th oldest divergence, where *n* is the number of sampled extant species *n* = ⎣*N*ρ⎤. FBD trees with ρ = 0.2, 0.5, and 0.8 are shown in (c), (d), and (e), respectively.

Here, we investigate the impacts of taxon-sampling density on Bayesian tip dating under the unresolved FBD process. We use synthetic data produced by simulations of nucleotide sequence evolution, fossil occurrence times, and diversified taxon sampling. Following the “branches-and-twigs” sampling strategy used by major genome-sequencing projects, we also reanalyse a published eutherian mammal data set. Our results provide insights into the factors affecting tip dating under the unresolved FBD model, with potential implications for the use of this approach in the phylogenomic era.

## Materials and Methods

### Simulation Study

#### Simulation of species trees

We generated complete species trees under the birth-death process using TreeSim (Stadler 2011) in R (R Core Team 2018). To match the general characteristics of phylogenomic data sets from groups such as eutherian mammals and modern birds, we used a constant speciation rate λ = 0.05 per Myr and extinction rate μ = 0.02 per Myr while conditioning on the root age (i.e., time of the most recent common ancestor, MRCA, of all sampled taxa) *t*_mrca_ = 100 Ma. After generating 1000 trees, we retained 100 trees with moderate numbers of extant taxa (50 ≤ *N* ≤ 100). These trees all have extant descendants on both sides of the root bifurcation, and so *t*_mrca_ is equivalent to the crown age *t*_c_ (i.e., time of the MRCA of all extant taxa). For comparison, we rescaled these trees to produce an additional set of trees with root age *t*_mrca_ = 300 Ma, following the approach used by Klopfstein et al. (2019). We note that across the 200 species trees, the effective rates of speciation and extinction vary around the values chosen for the initial simulations, given the properties of the birth-death process (Ricklefs 2007; Höhna et al. 2011) and the nature of our rejection sampling (Casella et al. 2004).

Based on the complete species trees, we sampled extant species using a diversified sampling strategy (Lambert and Stadler 2013; Zhang et al. 2016). With *N* extant species in the complete tree and a sampling fraction ρ, the number of sampled extant species is *n* = ⎣*N*ρ⎤(rounding to the nearest integer). These *n* species were chosen such that they were descended from the oldest *n* – 1 divergences in the reconstructed tree of extant species. This is equivalent to setting the cut-off time, *x*_cut_, to the (*n* – 1)th oldest divergence in the reconstructed tree and choosing only one representative species per clade after *x*_cut_. We set ρ to 0.1, 0.2, 0.5, 0.8, and 1.0 (i.e., complete sampling) to emulate an increasingly dense sampling process. This led to 5–10 (median 7), 10–20 (median 15), 25–50 (median 36.5), 40–80 (median 58.5), and 50–100 (median 73.5) extant species sampled across the 100 trees of *t*_mrca_ = 100 Ma (Fig. 1). Because the relative branch lengths were not changed during the tree rescaling described above, the sampled extant species are identical between the trees with *t*_mrca_ = 100 Ma and *t*_mrca_ = 300 Ma.

#### Simulation of fossil occurrences

Fossils were sampled along each of the complete species trees with a constant fossil recovery rate *ψ* = 0.003 (with reference to our previous study on the impacts of fossil occurrences; Luo et al. 2020), which models fossil occurrence as a continuous Poisson process (Heath et al. 2014). Because of the differences in branch lengths, this yielded 2–15 (median 7) fossil occurrences for trees of *t*_mrca_ = 100 Ma, compared with 6– 39 (median 21) fossil occurrences for those of *t*_mrca_ = 300 Ma. We ensured that at least one sampled fossil was older than *x*_cut_ when ρ = 0.1, to meet the assumptions of the diversified FBD process (Ronquist et al. 2016; Zhang et al. 2016). According to *x*_cut_ under each ρ value, we divided all sampled fossils into two categories: those sampled not later than *x*_cut_ (Category I; Fig. 1), and those sampled later than *x*_cut_ (Category II). After sampling the extant species (with ρ and *n* as above), keeping all sampled fossils or only those of Category I, and pruning all unsampled lineages, we had 1000 (100 × 5 × 2) FBD trees of *t*_mrca_ = 100 Ma as well as 1000 FBD trees of *t*_mrca_ = 300 Ma. It should be noted that among the 2000 FBD trees, there are 251 duplicates because Category II can be empty, especially under complete sampling of extant species (i.e., ρ = 1).

#### Simulation of nucleotide sequence evolution

Using the 100 complete species trees with *t*_mrca_= 100 Ma, we simulated the evolution of nucleotide sequences along the reconstructed trees of extant species only. We first assumed a strict clock with a rate of 10^-3^ substitutions/site/Myr. We then simulated sequence evolution using Seq-Gen v1.3.4 (Rambaut and Grassly 1997) to generate 100 sequence alignments, each composed of five ‘loci’ of length 1000 bp. To allow comparison with previous evaluations of tip dating, we matched the settings of Luo et al. (2020) for the relative substitution rates across loci, which were sampled from a Dirichlet distribution with *α* = 3. We also used an HKY+G substitution model with base frequencies {A:0.35, C:0.15, G:0.25, T:0.25}, transition/transversion ratio *κ* = 4.0, and gamma shape parameter of 0.5 with four rate categories. The resulting sequence alignments were directly used as sequences of the extant species sampled when ρ = 1, and were further pruned in accordance with those sampled under other ρ values as above. These procedures were repeatedly applied for trees of *t*_mrca_ = 300 Ma for simulating sequence evolution. We also applied a relaxed clock for reconstructed trees of *t*_mrca_ = 100 Ma, using the white-noise model of among-lineage rate variation with a mean of 10^-3^ substitutions/site/Myr and standard deviation of 2×10^-4^ substitutions/site/Myr in NELSI v0.2 (Lepage et al. 2007; Ho et al. 2015).

#### Settings for core analyses

Tip dating was first performed on synthetic data derived from the FBD trees of *t*_mrca_ = 100 Ma, comprising the fossil occurrence times of Category I and the nucleotide sequences of sampled extant species that were simulated under a strict clock. We used an FBD model with diversified sampling as the tree prior, assuming constant rates of speciation, extinction, and fossil recovery before *x*_cut_ and zero fossil sampling thereafter (Zhang et al. 2016). We assigned diffuse priors: beta(1,1) for both turnover *r* = μ / λ and fossil sampling proportion *s = ψ* / (*ψ* + μ), and exponential(10) for diversification rate *d* = λ − μ, which are reparameterizations of speciation rate λ, extinction rate μ, and fossil recovery rate *ψ* (Heath et al. 2014). The sampling fraction of extant species ρ was fixed to its true value. A strict-clock model was shared by the five loci, with a uniform(10^-6^,1) prior for the evolutionary rate. We applied an HKY+G model with four rate categories to each locus.

Fossil occurrence times were directly used as point values, and the diversified FBD process was conditioned on sampling at least one species and the root age *t*_mrca_. For the prior on *t*_mrca_, we either used a normal distribution with a mean matched to the true value and a standard deviation of 10 Myr, or specified a uniform(0,200) distribution. We also used normal(120,10) and normal(80,10) distributions, to evaluate the potential impacts of the *t*_mrca_ prior. A total of 2000 data sets are involved in the analyses described here, which we refer to as our “core” analyses (Table 1).

**Table 1.** Settings and scenarios explored in our simulation study.

| Analysis | Condition | True<br>$t_{\text{mrca}}$ | Clock model | $t_{\text{mrca}}$ prior | Tree<br>topology | No. of<br>data sets |
| --- | --- | --- | --- | --- | --- | --- |
| Core analyses | Fossils of Category I, diversified sampling, matched $\rho = 0.1, 0.2, 0.5, 0.8, 1$ | 100 | Strict–Strict | Normal(100,10) | Estimated | 500 |
|  |  | 100 | Strict–Strict | Uniform(0,200) | Estimated | 500 |
|  |  | 100 | Strict–Strict | Normal(120,10) | Estimated | 500 |
|  |  | 100 | Strict–Strict | Normal(80,10) | Estimated | 500 |
| Alternative analyses for comparison | Fossils of Category I, diversified sampling, matched $\rho = 0.1, 0.2, 0.5, 0.8, 1$ | 100 | Strict–Strict | Normal(100,10) | Fixed | 500 |
|  |  | 100 | Strict–Strict | Uniform(0,200) | Fixed | 500 |
|  |  | 100 | Strict–Strict | Normal(120,10) | Fixed | 500 |
|  |  | 100 | Strict–Strict | Normal(80,10) | Fixed | 500 |
|  |  | 300 | Strict–Strict | Normal(300,10) | Estimated | 500 |
|  |  | 100 | Strict–Relaxed | Normal(100,10) | Estimated | 500 |
|  |  | 100 | Relaxed–Relaxed | Normal(100,10) | Estimated | 500 |
| Analyses with model mismatches | Fossils of Category I and II | 100 | Strict–Strict | Normal(100,10) | Estimated | 358 |
|  | Random sampling | 100 | Strict–Strict | Normal(100,10) | Estimated | 500 |
|  | Fossils of Category I and II, random sampling | 100 | Strict–Strict | Normal(100,10) | Estimated | 358 |
| | Mismatched $\rho = 0.5$ | 100 | Strict–Strict | Normal(100,10) | Estimated | 400 |

Note: $t_{\text{mrca}}$ = root age, that is, time of the most recent common ancestor of all extant and extinct species; fossils of Category I = fossils not younger than the cut-off time, which depends on the number of sampled extant species; fossils of Category II = fossils younger than the cut-off time. The column “Clock model” gives the clock model used for simulation and that used for analysis.

#### Settings for alternative analyses for comparison

The core analyses typically co-estimated node times and tree topology. For comparison, we carried out further analyses of the same data using different settings. Under the normal(100,10), uniform(0,200), normal(120,10), and normal(80,10) *t*_mrca_ priors, we fixed tree topologies to match the FBD trees used for simulation, so that only node times of the trees were estimated. Other settings for these 2000 data sets followed those in the core analyses. We also applied the settings in the core analyses to data derived from trees of *t*_mrca_ = 300 Ma, except for a normal(300,10) prior on the root age *t*_mrca_.

To examine the impacts of rate variation among lineages, we used an uncorrelated lognormal relaxed clock to analyse the data simulated under a strict clock in our core analyses, with a uniform(10^-6^,1) prior for the mean clock rate and a normal(100,10) prior for *t*_mrca_. We also applied the same relaxed-clock model to nucleotide sequences simulated under a relaxed clock for trees of *t*_mrca_ = 100 Ma. The rest of the settings followed those in the core analyses. These two scenarios contributed a further 1000 data sets to our alternative analyses (Table 1).

#### Settings for analyses with model mismatches

We also explored three situations involving mismatched models of species sampling (Table 1), using nucleotide sequences generated under a strict clock and fossil occurrences derived from trees of *t*_mrca_ = 100 Ma. Given that true *x*_cut_ is generally unknown for real data, we first examined a situation where all fossil occurrences were included while presuming diversified sampling of extant taxa. In this case, we kept only data sets with non-empty fossil occurrences in Category II (358 out of 500). Second, when fossils of Category I (or both Category I and II) were involved, we adopted an FBD process that assumed random sampling rather than diversified sampling of extant species, with the sampling fraction of extant species ρ fixed to its true value. Third, because the true value of ρ is not always known, we fixed ρ to 0.5 under diversified sampling so that it is misspecified for most of the data sets. The other settings for each of the three situations were the same as in our core analyses under the normal(100,10) *t*_mrca_ prior. These three situations involved 358, 858 (500 + 358), and 400 data sets.

#### Data analyses

We ran tip-dating analyses using the BDSKY v1.4.5 package (Stadler et al. 2013) in BEAST v2.6 (Bouckaert et al. 2019), with posterior distributions estimated by Markov chain Monte Carlo (MCMC) sampling. The placement of each fossil was constrained so that it descended from its parent node in the FBD trees, thus allowing each to be resolved in a crown or stem position among its extant relative(s). We ran MCMC sampling in duplicate, with samples drawn every 2000–5000 steps over 40–100 million steps, depending on the data set, and with a discarded burn-in fraction of 0.25. We used Tracer v1.7 (Rambaut et al. 2018) to check that the effective sample sizes of parameters for the combined MCMC samples were greater than 100. We then used TreeAnnotator v2.6.0 to identify the maximum-clade-credibility tree among the sampled trees, with fossil taxa either retained or pruned using FullToExtantTreeConverter. With each of the 100 maximum-clade-credibility trees treated as an independent replicate under each scenario (Table 1), we examined the estimates of evolutionary parameters across our various scenarios, while highlighting the impacts of increasing sampling densities represented by ρ values.

Our conditioning on sampling at least one species in the inferences did not necessarily enforce extant descendants on both sides of the root bifurcation. In other words, a crown fossil could be placed on the stem lineage outside the clade containing the extant taxa. Since the stem fossil did not carry any morphological information to inform the root age, we focused on divergence times estimated for the crown age *t*_c_ to compare with the true root age *t*_mrca_ (which always has *t*_c_ = *t*_mrca_ during simulation). Posterior tree length was used to summarize the global node times in the maximum-clade-credibility tree without fossils, with either its standard form (sum of branch lengths) or its variants (sum of internal branch lengths, or sum of branch lengths except those of the two branches descending from the root). We also examined total evolutionary change (the product of clock rate and tree length) and clock rate because these parameters are relevant to the inference of divergence times. Estimates of tree topologies were evaluated by topological distance between the maximum-clade-credibility tree and the true topology using the Robinson-Foulds metric (Robinson and Foulds 1981) in the R package ape (Paradis et al. 2004; Popescu et al. 2012), whether or not the fossil taxa were pruned.

### Analyses of Empirical Data

#### Data set

To explore the effects of taxon-sampling density in phylogenomic data, we reanalysed a data set of eutherian mammals from a study by Ronquist et al. (2016), which was originally assembled by O’Leary et al. (2013). The data set includes 41 extant species, representing the four eutherian superorders (Xenarthra, Afrotheria, Laurasiatheria, and Euarchontoglires) and all living eutherian orders, as well as 33 eutherian or potentially eutherian fossils ranging from the 35.3-Myr-old *Leptictis dakotensis* to the 123.3-Myr-old *Eomaia scansoria*. Following Ronquist et al. (2016), we used point values for the ages of the fossils. The data set comprises 36,860 nucleotide sites from 22 nuclear protein-coding genes and five nuclear untranslated regions, along with 4541 discrete morphological characters.

#### Total-evidence tip dating

We first performed total-evidence tip dating with both the molecular sequences and morphological characters, as done by Ronquist et al. (2016) (Fig. 2). While accounting for diversified sampling of extant species, we conditioned the FBD process on the origin time *t*_o_ of eutherians using an offset-exponential prior with a mean 145 Ma and an offset 100 Ma. We specified lognormal(-2.5,0.7) priors for λ and μ, with a waiting time of 10 Myr for a cladogenetic event, and a lognormal(-5,0.8) prior for *ψ* with a waiting time of 100 Myr for a fossil occurrence. Variable morphological characters were analysed using the Mkv+G model. The molecular sequence data were partitioned into four subsets (three corresponding to the codon positions of the 22 protein-coding genes, and one comprising the five untranslated nuclear regions), with each subset assigned a separate GTR+G model. Following Ronquist et al. (2016), we applied a single uncorrelated lognormal relaxed-clock model to the morphological and molecular data, with a lognormal(-6.0,0.5) prior (expectation ∼2.5×10^-3^ substitutions per site per Myr) for the mean rate and an exponential(1.0) prior for the standard deviation. The extant species sampling fraction ρ was fixed to 0.01. For our three independent runs of MCMC sampling using the BDSKY package in BEAST v2.6, samples were drawn every 10,000 steps over a total of 400 million steps, with a burn-in fraction of 0.25. To address problems with rooting and MCMC mixing, we enforced a monophyly constraint on Boreoeutheria (Euarchontoglires + Laurasiatheria), with a normal(80,8) prior for its pan-group age (Ronquist et al. 2016).

**Figure 2.**
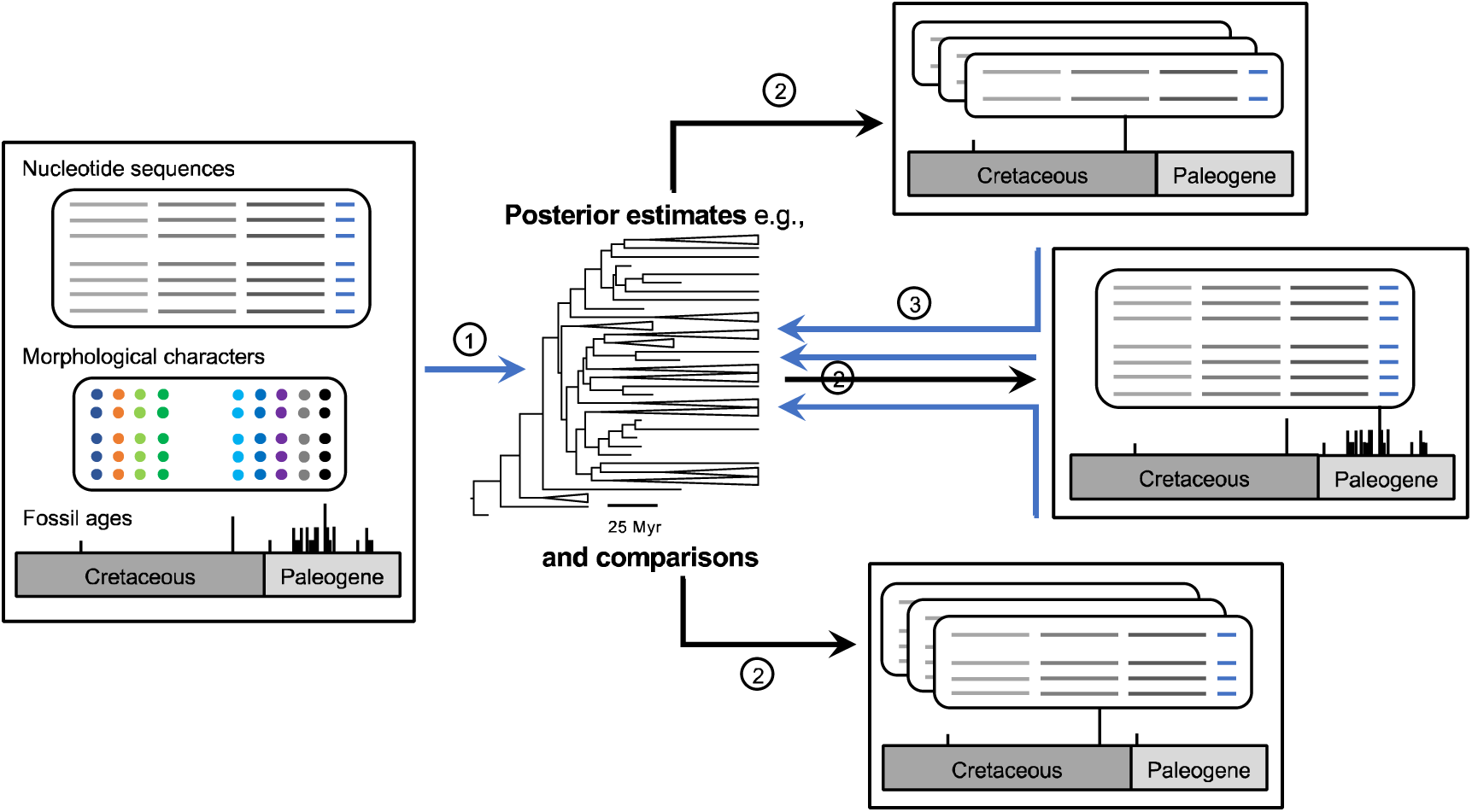
Flowchart showing the analysis pipeline in our empirical study of eutherian mammals. The steps are described in detail in the Materials and Methods. Briefly, we (1) performed total-evidence tip dating using molecular sequences from extant species and morphological characters from both extant species and 33 eutherian fossils. (2) Based on the posterior maximum-clade-credibility tree, representatives of extant species at the superorder, order, and species levels were then sampled, along with selected fossil occurrences. (3) Using molecular sequences of sampled extant species and fossil occurrences times, we carried out tip-dating analyses under the unresolved FBD process and compared the posterior estimates. Fossil occurrences are indicated by vertical bars along the geological timescale, with different heights denoting 1, 2, 3, and 4 occurrences at point ages. The figure indicates that 4, 5, and 33 fossil occurrences were selected at the superorder, order, and species levels, respectively, but we also included 28 fossil occurrences for analyses at the three levels (see Materials and Methods). Furthermore, we only show 3 (instead of 10 in practice) repeats of sampled extant species at higher levels.

#### Taxon sampling

Based on the maximum-clade-credibility tree from the total-evidence tip dating described above, we performed diversified sampling of extant species using a hierarchical approach similar to those used in major phylogenomic initiatives. To construct a superorder-level data set, we chose two representative species from each of the three superorders Afrotheria, Laurasiatheria, and Euarchontoglires, and one species from Xenarthra, with 10 random repeats. To construct 10 replicates of order-level data sets, we randomly selected a representative of each order of eutherian mammals. For each replicate at the superorder and order levels, we ensured that the MRCA of the chosen species from each group was as old as possible. Finally, we constructed a “species”-level data set comprising all extant species in the maximum-clade-credibility tree.

According to *x*_cut_ relative to the posterior medians of divergence times in the maximum-clade-credibility tree and considering fossil occurrences not younger than *x*_cut_, we first chose four (*Eomaia scansoria* at 123.3 Ma, *Maelestes gobiensis* at 77.8 Ma, *Ukhaatherium nessovi* at 77.8 Ma, and *Zalambdalestes lechei* at 77.8 Ma), five (adding *Protungulatum donnae* at 64.6 Ma), and all 33 fossil occurrence times at the superorder, order, and species levels, respectively. Second, in light of our simulation conditions (i.e., no stem fossil occurrence) and the first scenario explored in our analyses with model mismatches (i.e., fossils of Category I and II, diversified sampling), we eliminated the five stem fossil occurrences (*Eomaia scansoria*, *Leptictis dakotensis*, *Maelestes gobiensis*, *Ukhaatherium nessovi*, and *Zalambdalestes lechei*) in the maximum-clade-credibility tree, and used the remaining 28 fossil occurrence times at the three sampling levels.

#### Tip dating under the unresolved FBD process

Having molecular sequences for the sampled extant species and fossil occurrence times (i.e., without morphological characters), we carried out tip dating under the unresolved FBD process, with other settings and priors as in our total-evidence tip dating. We drew samples every 10,000 MCMC steps over a total of 300 to 400 million steps, with the first 25% discarded as burn-in. During MCMC sampling, we either put monophyly constraints on the fossil placements, or fixed the topology of all sampled taxa based on the maximum-clade-credibility tree derived from our total-evidence tip-dating analysis. While using posterior medians directly for analyses at the species level, we recorded medians across the 10 repeats for analyses at the superorder and order levels, respectively.

## Results

### Core Analyses

In our analyses of synthetic data without among-lineage rate variation, we found that increasing sampling density did not substantially improve estimates of divergence times. When we used a normal(100,10) root-age prior, which concentrates the prior density around the true value of t_mrca_, the crown age t_c_ was invariably estimated to be equal to estimated t_mrca_, as expected. Hereafter, we refer only to estimates of t_c_ except where otherwise noted. Estimates of t_c_ was recovered accurately regardless of whether the sampling fraction ρ was 0.1 or 1 (Fig. 3a and 3b). Specifically, medians of the 100 replicates of the posterior medians for t_c_ were 101, 103, 104, 104, and 104 Ma, while those of the 95% credibility interval (CI) were 89–116, 91–118, 93–118, 92–118, and 93–118 Ma when ρ = 0.1, 0.2, 0.5, 0.8, and 1, respectively. Standardized tree-length estimates approached the expected values and were similar across different values of ρ, though with slight variation in 95% CIs. Excluding the two branches descending from the root did not alter these patterns, except for a slightly larger variation across the repeats when ρ = 0.1. However, there was an increase in precision with increasing ρ when the terminal branches were excluded. In general, standardized total evolutionary change, using the substitution rate measured by the clockRate parameter, did not vary much with ρ; however, medians were less variable across replicates and 95% CIs were slightly wider when ρ was larger.

**Figure 3.**
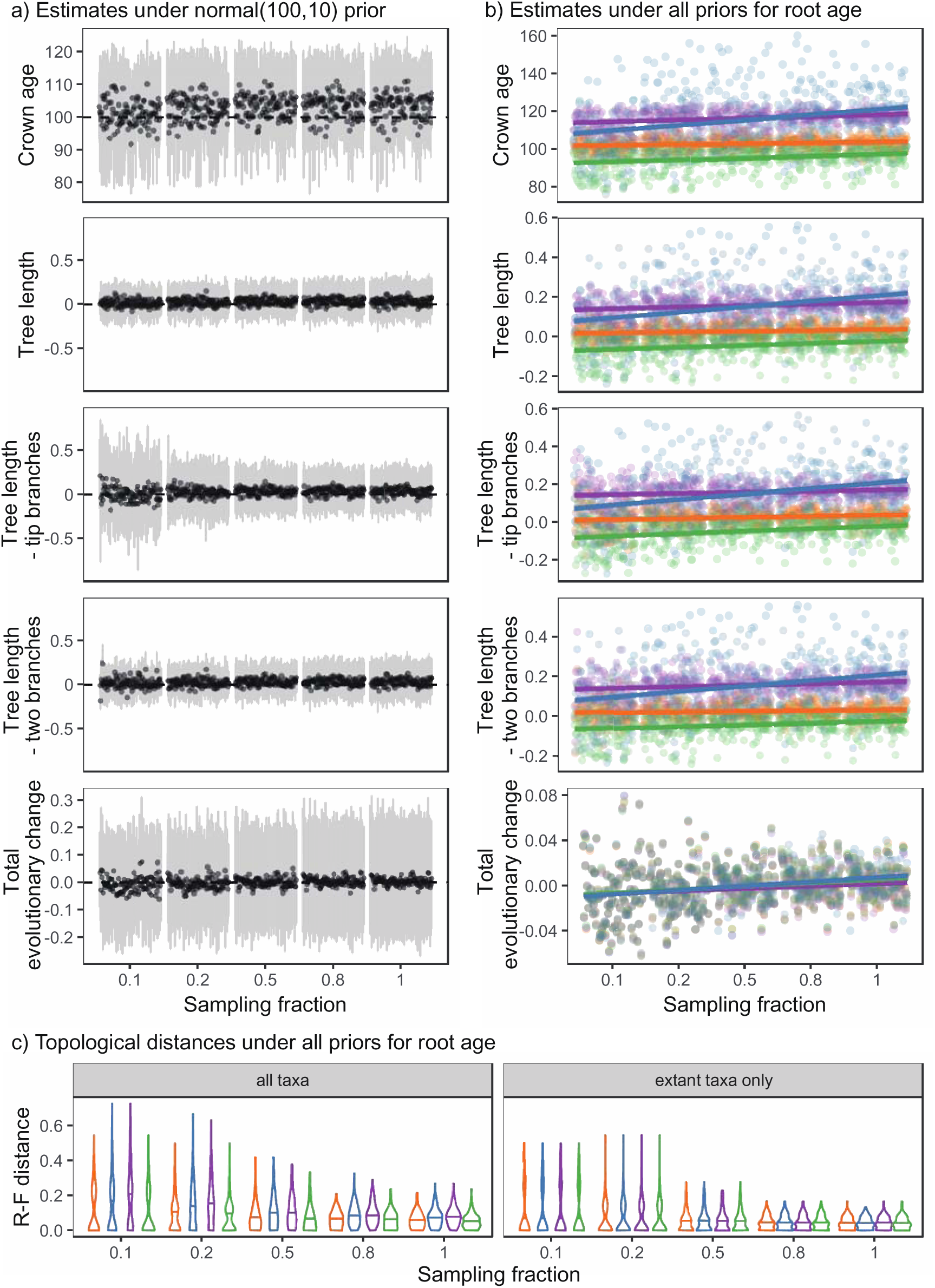
Posterior estimates from tip dating under the unresolved FBD process for time and topology in our core analyses. (a) Time estimates under a normal(100,10) root-age *t*_mrca_ prior, with black circles representing posterior medians and gray lines representing 95% credibility intervals (CIs). For crown age *t*_c_, absolute estimates are shown. For tree length (i.e., sum of all branch lengths), tree length without tip branches (i.e., sum of internal branch lengths), tree length without two branches (i.e., sum of all branch lengths except those of the two branches descending from the root), and total divergence (i.e., the product of clock rate and tree length) from the top two to the bottom, standardized estimates (i.e., distances between the estimates and the true values, divided by the true values) are shown. There are 100 replicates at each level of sampling fraction (*x*-axis). Dashed horizontal lines indicate the true or expected values. (b) Time estimates under normal(100,10), uniform(0,200), normal(120,10) and normal(80,10) priors for *t*_mrca_, denoted by orange, blue, purple, and green, respectively. For the sake of clarity, only absolute or standardized posterior medians are shown, together with lines denoting linear-regression results (by R package ggplot2; Wickham 2016). There are 100 replicates at each level of sampling fraction under each *t*_mrca_ prior. (c) Violin-plot summaries of topological distances under normal(100,10), uniform(0,200), normal(120,10) and normal(80,10) priors for *t*_mrca_, denoted by orange, blue, purple, and green, respectively. Each summary is based on 100 replicates, with a horizontal line indicating the median. The left panel shows the corrected Robinson-Foulds (R-F) distances between maximum-clade-credibility trees and true trees with fossils included at each level of sampling fraction, whereas the right panel shows those with fossils pruned.

To some extent, increasing sampling density improved the inference of tree topologies for analyses under the normal(100,10) *t*_mrca_ prior, as topological distances (measured by the Robinson-Foulds metric corrected by the number of tips) tended to be less variable with smaller medians across replicates at higher sampling densities (Fig. 3c). When fossils were pruned from the trees, the accuracy was higher than when the fossils were included for comparison. This is due to the fact that only fossil occurrence times were used and their phylogenetic positions were mainly informed by topological constraints and the FBD prior.

### Impacts of Altering the Root-Age Prior

Altering the prior on t_mrca_ had noticeable impacts on the time estimates (Fig. 3b). As expected, using normal(120,10) and normal(80,10) priors led to over- and underestimates, respectively, of crown age t_c_ and global node times. Using a uniform(0,200) prior also produced overestimated node times; the effect was similar to using a normal(120,10) prior but the posterior medians were more variable. Nevertheless, the estimates of total evolutionary change were relatively robust to the choice of root-age prior, with medians of the standardized metric around zero.

Increasing sampling density had only small impacts on the estimates of node times. Under the normal(120,10) *t*_mrca_ prior, posterior medians were less variable with larger ρ. For example, standard deviations for *t*_c_ declined from 5.2 Ma for ρ = 0.1 to 2.5 Ma for ρ = 1. Under the uniform(0,200) *t*_mrca_ prior, posterior node times increased and became more uncertain with ρ; the 95% CIs for *t*_c_ were 89–124 and 93–174 for ρ = 0.1 and 1, respectively. Increasing ρ did not produce other obvious effects under the normal(80,10) *t*_mrca_ prior, except for a reduced variation in estimated tree length when excluding terminal branches. We note that with reference to true values, increasing sampling density had a negative impact under the normal(120,10) and uniform(0,200) *t*_mrca_ priors, but a slightly positive impact under the normal(80,10) prior.

Topological distances under the three alternative *t*_mrca_ priors were similar to those under the normal(100,10) prior, especially for tree topologies of only extant tips (Fig. 3c). Yet, the uniform(0,200) and normal(120,10) priors led to poorer topological inferences when fossils were included, with topological distances declining as ρ increased.

### Impacts of Fixing the Tree Topology

Based on the patterns of estimates being consistent for key node times with those for tree length from our alternative analyses for comparison, we further focused on *t*_c_ with relative metrics only. We found that fixing tree topologies to those used for simulation, as well as increasing sampling density, generally improved both accuracy and precision of estimates (Fig. 4). Under the normal(100,10) and normal(80,10) *t*_mrca_ priors, the relative 95% CI width for *t*_c_ became smaller as ρ increased. The relative bias also improved slightly under the normal(100,10) prior. Under the normal(120,10) prior, both the relative bias and relative 95% CI width approached the expected values when ρ became larger. The positive effects of increasing ρ were most prominent under the uniform(0,200) prior; estimates of *t*_c_ had much better accuracy and precision, in contrast with the tendency of further deviation with larger ρ in the core analyses. Relative to results from the normal(100,10) prior, however, analyses with the normal(120,10) and normal(80,10) priors still produced over- and underestimations, while estimates were more variable under the uniform(0,200) prior.

**Figure 4.**
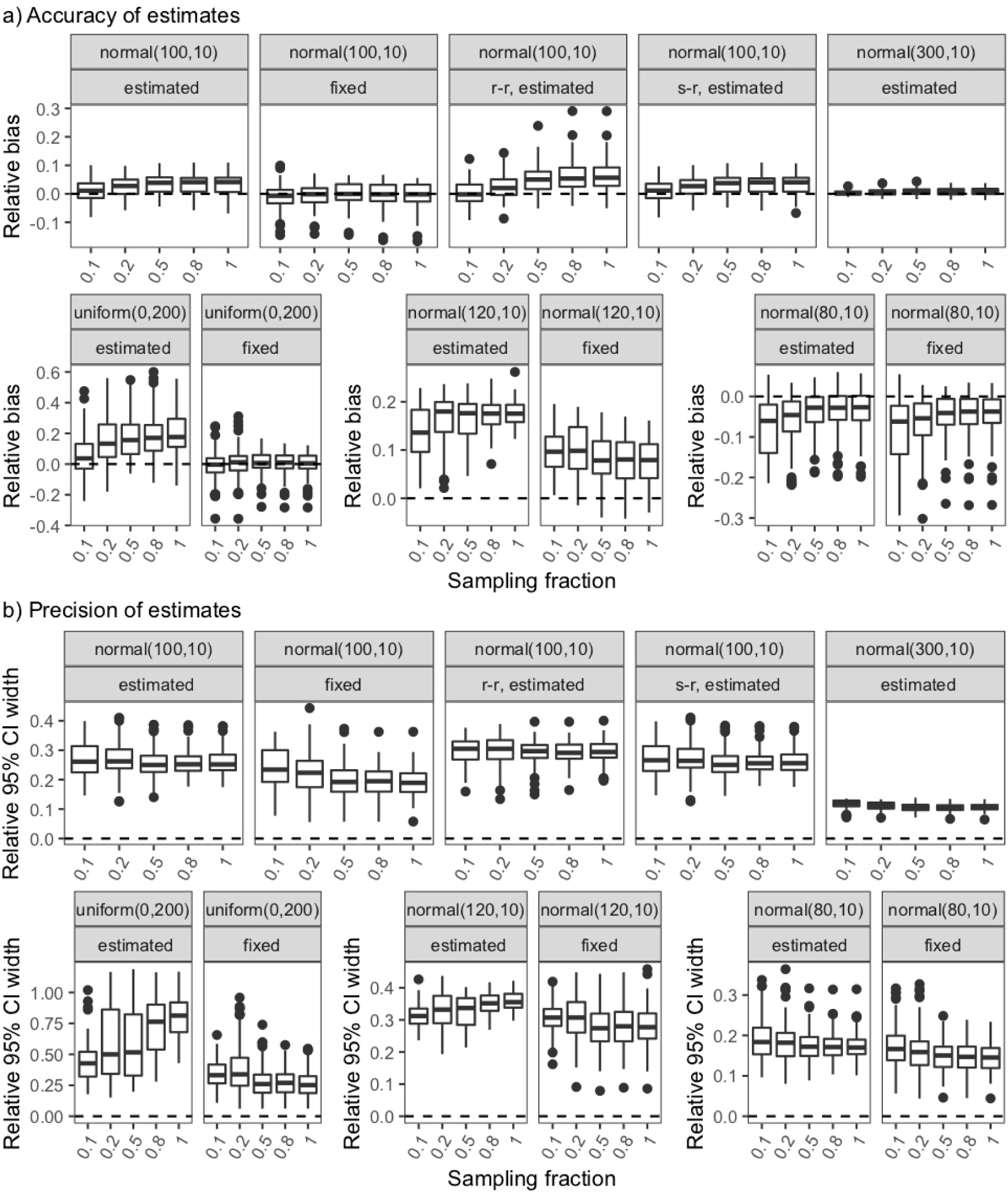
Posterior time estimates from tip dating under the unresolved FBD process in our core analyses and alternative analyses for comparison. (a) Boxplots to summarize accuracy of estimates for crown age *t*_c_, as measured by relative bias (distance between the posterior median and the true value, divided by the true value). (b) Boxplots to summarize precision of estimates for *t*_c_, as measured by relative 95% credibility interval width (the posterior 95% CI width divided by the true value). For both (a) and (b), each combination of panels shows comparable results from different settings. The upper combination from left to right: a normal(100,10) *t*_mrca_ prior with tree topologies estimated, ditto with topologies fixed, ditto with topologies estimated and a relaxed clock applied to data simulated using a relaxed clock (denoted by “r-r”), ditto with topologies estimated and a relaxed clock applied to data simulated using a strict clock (denoted by “s-r”), and a normal(300,10) *t*_mrca_ prior with topologies estimated; the bottom left combination: a uniform(0,200) *t*_mrca_ prior with topologies estimated and ditto with topologies fixed; the bottom middle combination: a normal(120,10) *t*_mrca_ prior with topologies estimated and ditto with topologies fixed; the bottom right combination: a normal(80,10) *t*_mrca_ prior with topologies estimated and ditto with topologies fixed. Dashed horizontal lines indicate the true or expected value for each summary, which is based on 100 replicates at each level of sampling fraction.

Posterior medians of clock rates were estimated to be inversely proportional to relative biases for *t*_c_, regardless of whether the tree topologies were fixed (in our alternative analyses) or not (in our core analyses) (Fig. 5a). For example, under the normal(100,10) *t*_mrca_ prior, the relative biases for *t*_c_ were around 0.0, and the medians for clock rates approached the expected value of 0.001. Under the normal(120,10) *t*_mrca_ prior and with tree topologies fixed, posterior median clock rates increased with ρ, while relative biases for *t*_c_ decreased. This relationship was generally consistent with our estimates of the total evolutionary change, which is a product of tree length and clock rate.

**Figure 5.**
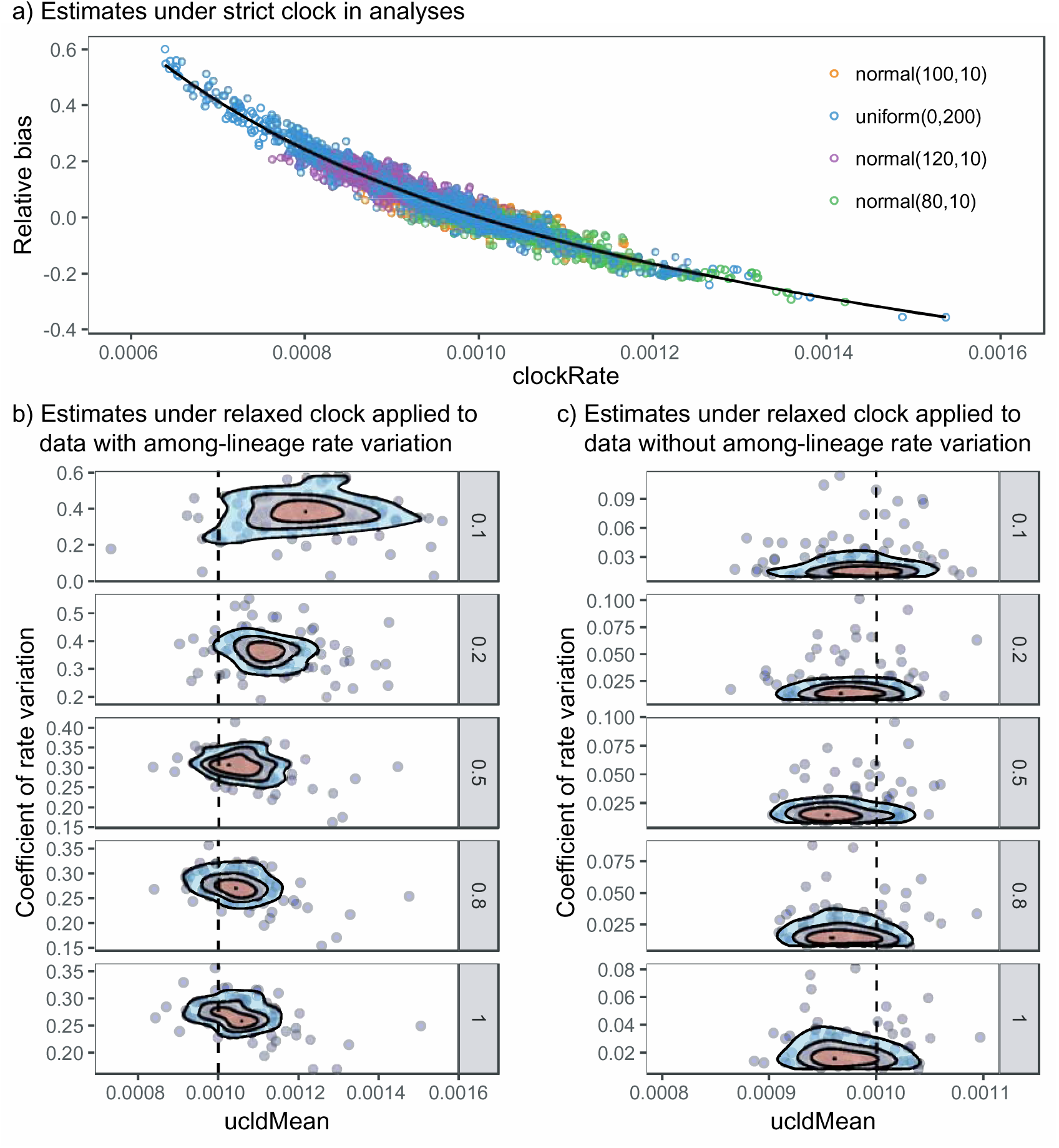
Posterior rate estimates from tip dating under the unresolved FBD process in our core analyses and alternative analyses for comparison. (a) Scatterplot for posterior medians of clock rates (measured by clockRate) against relative biases of time estimates resulted from analyses employing a strict clock. Estimates under normal(100,10), uniform(0,200), normal(120,10) and normal(80,10) priors for *t*_mrca_ are shown by orange, blue, purple, and green, respectively, while overlooking whether tree topologies were fixed or not during analyses. The black curve derives from a polynomial-regression analysis (by ggplot2). (b) Density scatterplots for posterior medians of mean clock rates among lineages (measured by ucldMean) against those of rate variation among lineages (measured by coefficientOfVariation) for analyses with a relaxed clock applied to data simulated using a relaxed clock. (c) Ditto for analyses with a relaxed clock applied to data simulated using a strict clock. For (b) and (c), at each level of sampling fraction, increasingly orange shades indicate higher density among the 100 replicates; dashed vertical lines indicate the expected value of mean clock rate among lineages.

### Impacts of Deep Divergence or Among-Lineage Rate Variation

Our alternative analyses also tested the effects of increasing sampling density on data derived from FBD trees of *t*_mrca_ = 300 Ma. We found that analyses with a normal(300,10) *t*_mrca_ prior produced accurate and precise estimates of *t*_c_ for all values of ρ (Fig. 4). Clock rates were also well estimated by posterior medians, approaching the true value of 0.001.

When we applied a relaxed-clock model to data with among-lineage rate variation, we found that *t*_c_ was more clearly overestimated with larger ρ, and the relative 95% CI widths were greater than those from comparable data in our core analyses (Fig. 4). On the other hand, estimated clock rates (measured by posterior medians for the ucldMean parameter) decreased and became more accurate with increasing ρ (Fig. 5b). As a result, in contrast with the results from the data sets described above, total evolutionary change was also better estimated when ρ was larger. When we applied the relaxed-clock model to data without among-lineage rate variation, however, estimates of *t*_c_ and clock rates were similar to those from comparable data in our core analyses (Fig. 4 and 5c).

With fossils included, corrected Robinson-Foulds distances were slightly larger for analyses involving deep divergence than for our core analyses under the normal(100,10) *t*_mrca_ prior or those involving among-lineage rate variation. This was consistent with larger numbers of fossils being included in analyses under the normal(300,10) *t*_mrca_ prior. With fossils excluded, there were no obvious differences in topological distances among these analyses. A relaxed clock applied to data without among-lineage rate variation did not lead to substantially different topological inferences compared with our core analyses under the normal(100,10) *t*_mrca_ prior.

### Impacts of Mismatched Species-Sampling Models

Our final analyses of synthetic data involved mismatched models of species sampling, which generally resulted in posterior medians of *t*_c_ near the expected value (Fig. 6). Compared with the results from comparable data in our core analyses (i.e., under the normal(100,10) *t*_mrca_ prior; the control), when we used a diversified sampling model to analyse data sets that included all fossils, *t*_c_ was obviously underestimated. However, when random sampling of extant species was assumed, *t*_c_ was overestimated for lower values of ρ, and appeared no different from that of the control for ρ = 1. When all fossils were included and random sampling was assumed, estimates of *t*_c_ were intermediate between those under the first two scenarios, and overestimation occurred only with smaller ρ. When ρ was fixed to 0.5 for data with actual ρ > 0.5, *t*_c_ was slightly overestimated; however, there was little impact when ρ was incorrectly fixed to 0.5 for data with ρ < 0.5. Except for the scenario of mismatched ρ the posterior medians of *t*_c_ across these scenarios were expected to reach similar values at ρ = 1, but following different trajectories at different values of ρ. For instance, relative biases of *t*_c_ roughly increased with ρ when we used a diversified sampling model to analyse data sets that included all fossils.

**Figure 6.**
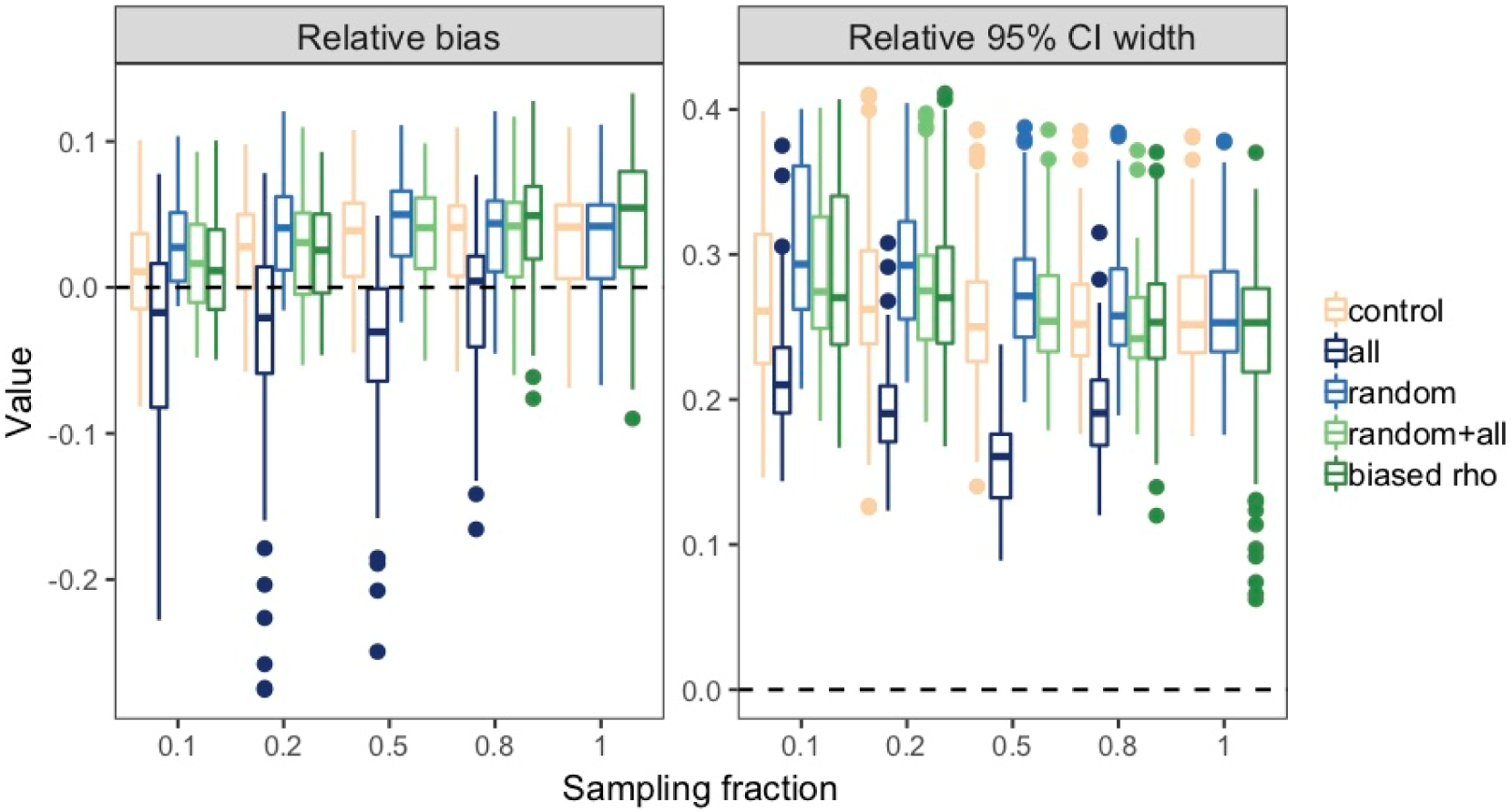
Posterior estimates from tip dating under the unresolved FBD process for crown age *t*_c_ in our analyses with model mismatches. Accuracy and precision were measured by relative bias and relative 95% confidence interval (CI) width, respectively. At each level of sampling fraction, results are shown for: counterpart in our core analyses (denoted by “control”); those with all fossils included (denoted by “all”); those with random sampling assumed (denoted by “random”); those with random sampling assumed and all fossils included (denoted by “random+all”); and those with mismatching sampling fraction (denoted by “biased rho”) from left to right (where appropriate). Boxplot summaries are based on varying numbers of replicates (Table 1). Dashed horizontal lines indicate expected values.

Among these analyses, including all fossils had different impacts on the precision of *t*_c_ estimates, in that the relative 95% CI widths deviated less from the expected value of zero and had lower variance across replicates. In contrast, when random sampling was assumed or ρ was fixed to a value larger than the true value, the relative 95% CI widths were further from the expected value (Fig. 6). With reference to the control, ρ mismatches had limited impacts on the precision of estimated *t*_c_. Among these analyses, precision of *t*_c_ improved or remained roughly constant when true ρ increased.

Similar to the above analyses using data without among-lineage rate variation, with these model mismatches, estimates of clock rates under a strict-clock model were inversely proportional to posterior medians or relative biases of *t*_c_. These results point to good estimation of total evolutionary change. Model mismatches had little impact on estimates of tree topologies, with the exception that including all fossils led to greater or more variable topological distances from the true trees when ρ was small.

### Analysis of Eutherian Mammals

Our total-evidence tip-dating analysis of eutherian mammals produced estimates of relationships that are consistent with previous inferences (Fig. 7a versus figure 8 in Ronquist et al. 2016). These include monophyly of each of the four mammalian superorders and monophyly of Euarchonta (primates with tree shrews and flying lemurs). Posterior probabilities for most groupings are greater than 75%. However, our tree also shows some differences from the previous estimate by Ronquist et al. (2016). For extant taxa, we found a sister relationship between Caviidae (*Cavia*) and Sciuridae (*Ictidomys*) in rodents, rather than a grouping of Castoridae (*Castor*), Muridae (*Rattus*), and Caviidae (*Cavia*). Our tree resolved some polytomies involving fossil taxa such as *Hyopsodus paulus*. The placements of some fossil taxa also differ, such as the position of *Sinopa rapax* in crown Carnivora rather than on the stem lineage.

**Figure 7.**
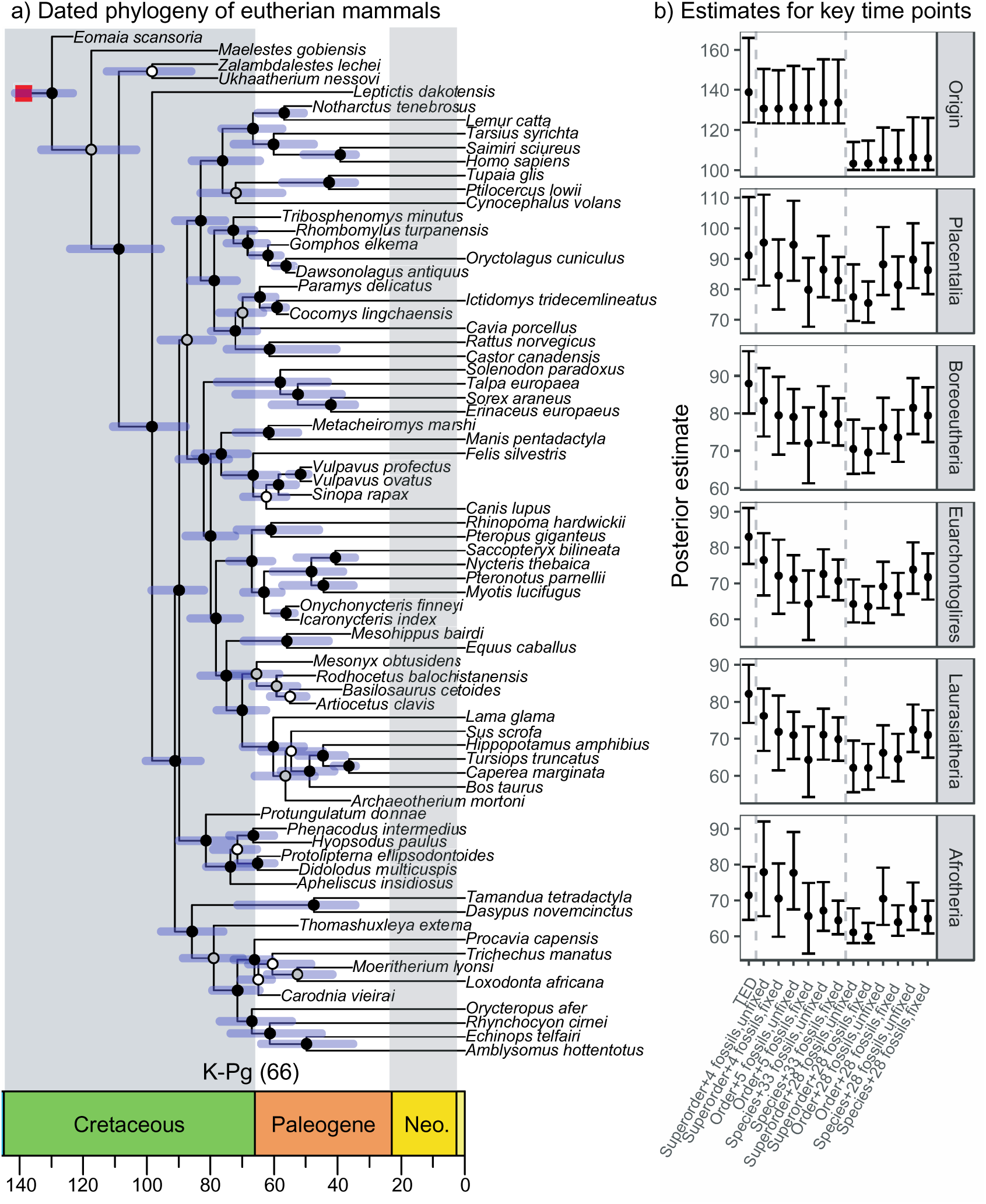
(a) Posterior maximum-clade-credibility tree of eutherian mammals from our total-evidence tip dating. Timeline at the bottom of the tree is in accordance with posterior medians of divergence times, and blue horizontal bars in the tree indicate 95% credibility intervals. Colored circles represent posterior probabilities: ≥ 95% (black), <95% but ≥75% (gray), and < 75% (white). The red square denotes the origin time of eutherian mammals, without 95% credibility interval. (b) Posterior estimates for the origin time of eutherian mammals, crown age of Placentalia, and node times for Boreoeutheria, Euarchontoglire, Laurasiatheria, and Afrotheria in the maximum-clade-credibility trees with fossils pruned. Absolute values are shown, with circles representing posterior medians and lines representing 95% credibility intervals. For each panel from left to right, results are from: total-evidence tip dating (denoted by “TED”); tip dating with extant species sampled at the superorder level, four fossil occurrences, and tree topologies estimated (denoted by “Superorder+4 fossils, unfixed”); ditto with tree topologies fixed (denoted by “Superorder+4 fossils, fixed”); tip dating with extant species sampled at the order level, five fossil occurrences, and tree topologies estimated (denoted by “Order+5 fossils, unfixed”); ditto with tree topologies fixed (denoted by “Order+5 fossils, fixed”); tip dating with extant species sampled at the species level, 33 fossil occurrences, and tree topologies estimated (denoted by “Species+33 fossils, unfixed”); ditto with tree topologies fixed (denoted by “Species+33 fossils, fixed”); tip dating with extant species sampled at the superorder level, 28 fossil occurrences, and tree topologies estimated (denoted by “Superorder+28 fossils, unfixed”); ditto with tree topologies fixed (denoted by “Superorder+28 fossils, fixed”); tip dating with extant species sampled at the order level, 28 fossil occurrences, and tree topologies estimated (denoted by “Order+28 fossils, unfixed”); ditto with tree topologies fixed (denoted by “Order+28 fossils, fixed”); tip dating with extant species sampled at the species level, 28 fossil occurrences, and tree topologies estimated (denoted by “Species+28 fossils, unfixed”); and ditto with tree topologies fixed (denoted by “Species+28 fossils, fixed”). Vertical lines separate the results from our total-evidence tip dating, tip dating with fossils selected at the three levels, and tip dating with 28 fossils at the three levels.

To compare our estimates with those from other tip-dating analyses, we focused on time estimates for the origin, Placentalia, Boreoeutheria, Euarchontoglires, Laurasiatheria, and Afrotheria. Our estimates are generally younger than those under uninformative priors but older than those under informative priors by Ronquist et al. (2016). For example, our tree placed the placental crown radiation in the Cretaceous (median 91 Ma, 95% CI 83–100 Ma) and estimated that Boreoeutheria diverged into Euarchontoglires and Laurasiatheria about 87 Ma when fossils were pruned. Additionally, we found that all orders in Boreoeutheria diverged from each other before the Cretaceous-Paleogene mass extinction (66 Ma), whereas divergences among some Afrotheria orders occurred afterwards (Fig. 7a).

We then examined the impacts of taxon sampling on divergence-time estimates in the maximum-clade-credibility trees with fossils pruned. When fossil occurrences were also sampled at corresponding densities (i.e., 4, 5, and 33 fossils at superorder, order, and species levels, respectively), the effects on time estimates were mixed. However, except for the origin time, there was a tendency for posterior medians to provide younger ages for each node at each sampling level when topologies of all sampled taxa were fixed than when only constraints on fossil placements were imposed (Fig. 7b). In comparison, analyses including 28 fossils at each sampling level produced clearer patterns, where the posterior medians at the superorder level were obviously younger than those from the corresponding analyses with four sampled fossil occurrences. Posterior medians increased with sampling density regardless of whether tree topologies were fixed or not (with the exception of Afrotheria when tree topologies were not fixed), although the degree differed among nodes (e.g., the origin versus Placentalia). Fixing tree topologies led to younger estimates at each sampling level, except for the origin time. In addition, we found that sampling density had almost no effect on the estimate of the origin time. Instead, fossil occurrences (or even the stem fossil occurrences) seemed to be more important for the estimate of the origin time, but they had limited effects on other node times.

## Discussion

Following the design of major phylogenomic initiatives, we have examined the effects of taxon-sampling schemes on Bayesian tip dating under the unresolved FBD process. Phylogenetic studies have consistently striven for denser taxon sampling, which evidently has positive effects on estimates of tree topologies and branch lengths (e.g., Pollock et al. 2002; Hugall and Lee 2007; Heath et al. 2008). It is also an important consideration for Bayesian molecular dating (Bromham et al. 2018), given the relationship of taxon sampling to important factors such as tree imbalance (Duchêne et al. 2015), rate variation among lineages (Soares and Schrago 2015), and bare-branch attraction (Spasojevic et al. 2021).

Our simulation study has shown that increasing the density of taxon sampling does not necessarily lead to better estimates of divergence times, and that the dates of key nodes can potentially be estimated reasonably well even with relatively sparse taxon sampling. These results are expected, to some extent, in the phylogenomic era, given that a larger amount of sequence data for each taxon could offset some of the disadvantages of sparse taxon sampling (Hillis et al. 1994). Denser taxon sampling can mitigate the impacts of some phylogenetic artifacts (Kapli et al. 2020; as exemplified by Fig. 3c in this study), but other factors such as data type, model choice, analytical methods, and missing data can also be important (e.g., Peloso et al. 2016; Streicher et al. 2016; Reddy et al. 2017; Prasanna et al. 2020). Moreover, Bayesian tip dating can at least partly account for diversified taxon sampling during co-estimation of tree topology and divergence times, potentially reducing some of the negative impacts of poor taxon sampling (Heath et al. 2008, 2014; Zhang et al. 2016). Therefore, we recommend making use of these models in phylogenomic dating analyses.

### Interactions Between Taxon-Sampling Density and Other Factors

Our results reveal the interaction between taxon-sampling density and other factors in Bayesian tip dating. Most of our analyses used synthetic data that had been generated without among-lineage rate variation, which resulted in good estimates of the total evolutionary change, regardless of whether the sampling fraction of extant species ρ was 0.1 or 1, or whether the data were analysed using a strict-clock or relaxed-clock model. As expected, estimated divergence times and clock rate were inversely proportional to each other, while being subject to varying levels of taxon-sampling density and other factors (see below). However, in the presence of data with among-lineage rate variation, sparse taxon sampling led to overestimates of the clock rate and the amount of evolutionary change, although estimates of divergence times were still negatively correlated to those of clock rate. Thus, our results highlight a potential benefit to increasing the density of taxon sampling, even though divergence times were generally well estimated under all degrees of taxon sampling. This is consistent with previous findings of the benefits of denser taxon sampling in Bayesian molecular dating for data with among-lineage rate variation (Soares and Schrago 2015).

One appealing feature of Bayesian tip dating under the resolved FBD process is the capacity to resolve the phylogenetic placement of a fossil as a tip or sampled ancestor (i.e., a tip with zero-length terminal branch). Under the unresolved FBD process, however, the fossil placement is a stochastic variable conditioned on a topological constraint, and the fossil is usually pruned from the dated tree (e.g., Heath et al. 2014; O’Reilly and Donoghue 2020). To examine the potential influence of the topological constraints on fossil placements, we obtain maximum-clade-credibility trees with fossils included in addition to those with extant species only, and computed their topological distances from the true trees. Using data without among-lineage rate variation, we found that topological constraints helped to produce more accurate fossil positions as taxon-sampling density increased. However, there was not necessarily a positive relationship to accurate estimates of divergence times. When we fixed tree topologies to those used for simulation (i.e., *a priori* accurate positions of both extant and fossil species), estimates of node times generally improved with the density of taxon sampling, although the specific effects of denser taxon sampling appeared to vary with the root-age prior (see below).

In most of the treatments examined in our simulation study, we matched the models used for simulation and analysis. In reality, however, the evolutionary process is unobserved and phylogenetic models are invariably violated to some extent. When we analysed data that had been generated without among-lineage rate variation, we found that wrongly assuming random taxon sampling led to overestimates of divergence times under sparse taxon sampling, echoing the findings of a previous study that dated the early radiation of hymenopteran insects to the Triassic and Permian (∼252 Ma) under a model of diversified taxon sampling versus the Carboniferous (∼347 Ma) under a model of random taxon sampling (Zhang et al. 2016). For some groups of organisms such as fishes (Matschiner et al. 2017), fossil occurrence records are abundant, and fossil subsampling can be carried out (Matschiner 2019; O’Reilly and Donoghue 2019). In contrast, in tip-dating analyses of groups that have a more limited fossil record, it might be necessary to include even very young fossil occurrences. Our analyses involving fossils younger than the cut-off time *x*_cut_ produced underestimates of node times, particularly with sparse taxon sampling. Denser taxon sampling thus should be beneficial due to a reduction in model misspecification with increasing sampling. We note that our analyses with model mismatches did not produce very large discrepancies (especially under the other two scenarios), but these problems are likely to be worse in analyses of empirical data, where there is a much greater risk of model misspecification.

### Root-Age Prior for Bayesian Tip Dating

While fossil tips provide calibrating information, Bayesian tip dating usually needs the FBD tree prior to be conditioned on an informed starting point such as the origin time *t*_o_ or the root age *t*_mrca_ (Gavryushkina et al. 2014; Zhang et al. 2016). Our analyses of data without among-lineage rate variation have shown that the root-age prior has a much greater impact than taxon-sampling density on estimates of node times, as reflected in the time estimates under the three normal-distribution priors. When we specified root-age priors that placed high probability near the true values, evolutionary parameters tended to be accurately estimated and showed little variation across different densities of taxon sampling.

The impact of the root-age prior is not particularly surprising, given the substantial influences of such calibrations in node-dating approaches (Duchêne et al. 2014; Bromham 2019). Although both the unresolved and resolved FBD processes treat fossils as tips (Ronquist et al. 2012; Heath et al. 2014), tip dating under the former is more similar to node dating with regard to topological constraints for fossil affinities (Zhang et al. 2016). The root-age prior effectively functions as an additional calibration for tip dating under the unresolved FBD process, with potential impacts on posterior time estimates (Ho and Phillips 2009; Battistuzzi et al. 2015; Álvarez-Carretero et al. 2019). In our case, it also appeared to have a slight influence on posteriors of fossil placements, which could inflate biases in time and rate estimates. This potential effect is suggested by the improvements seen in our analyses with fixed tree topologies.

The lower bound of the uniform(0,200) prior for the root age was always limited by the oldest fossil occurrence, so the actual (or induced) mean was larger than the true root age of 100 Ma. When we used a uniform(0,200) *t*_mrca_ prior without fixed tree topologies, we obtained results that were consistent with those from a previous simulation study using molecular data under the unresolved FBD process, but different from the accurate results that were obtained using both molecular and morphological data under the resolved FBD process (Luo et al. 2020). This highlights the different effects of the root-age or origin-time prior on total-evidence tip dating and unresolved-FBD tip dating.

### Diversified Sampling Strategy and Taxonomical Ranks

Our exploration of different degrees of taxon sampling was inspired by the “branches-and-twigs” sampling strategies adopted by major genome-sequencing initiatives (e.g., Jarvis et al. 2014; Lewin et al. 2018; Fan et al. 2020). In the Fish10K initiative, for example, Phase I involves sequencing genomes of 500 representatives at the order level, Phase II targets 3500 species representing all families, and Phase III expands the sampling to 10,000 species that represent all genera (Fan et al. 2020). To mimic this strategy, we explored a series of values for the sampling fraction ρ, while sampling extant species using the diversified sampling strategy at each level of ρ (Höhna et al. 2011; Lambert and Stadler 2013; Zhang et al. 2016), as exemplified in Figure 1.

Under diversified sampling schemes (at each ρ < 1), the tree has long terminal branches that maximize the tree length of the chronogram of extant taxa. In effect, this tree balances the number of extant descendants on either side of each of the deep divergence events. For example, subject to the cut-off time *x*_cut_ in Figure 1b, it is possible that we actually sample five extant species that representing either five orders or two orders (with four species from one order and one species from the other order). In reality, taxa are usually sampled on the basis of their taxonomical ranks and other factors (e.g., specimen availability and a balance between various groups), and these sampled lineages would not necessarily be the same as those defined by a single cut-off time in the chronogram. For instance, our empirical analyses sampled two representatives from each of the three superorders Afrotheria, Laurasiatheria, and Euarchontoglires, and one representative from Xenarthra. This is not fully consistent with the seven exemplars sampled under the diversified sampling strategy based on the dated eutherian phylogeny in Figure 7a. Therefore, although the diversified sampling strategy can account for incomplete taxon sampling to a large extent (Donoghue and Yang 2016; Zhang et al. 2016), it cannot completely model the taxonomically “diversified” sampling schemes that are used in reality.

### Evolutionary Timescale of Eutherian Mammals

In our analysis of a phylogenomic data set from eutherian mammals, we found several patterns that were generally consistent with those from our simulation study. These included the influences of fixing the tree topology, and when all crown fossils were included, an increase in estimated node times with denser extant taxon sampling, and a decrease in estimated node times. However, we are unable to gauge the accuracy of these estimates.

The diversification history of eutherian mammals continues to be debated, with different sources of data or analytical methods resulting in various scenarios (e.g., dos Reis et al. 2012; O’Leary et al. 2013; Phillips 2016; Ronquist et al. 2016; Davies et al. 2017). For instance, O’Leary et al. (2013) found support for an explosive model of placental radiations around the Cretaceous-Paleogene boundary, whereas our estimates were more consistent with a fuse model (Phillips 2016). Our analyses highlighted the impacts of including morphological characters on inferences of eutherian evolutionary history; out total-evidence dating analysis yielded older estimates than tip dating under the unresolved FBD process at the species level, a result that is consistent with previous findings of potential date overestimation by total-evidence dating (e.g., Wood et al. 2013; O’Reilly et al. 2015; Ronquist et al. 2016). In addition, informative priors that placed penalties on ghost lineages had profound effects, helping to close the gap between paleontological and molecular evidence as shown by Ronquist et al. (2016). Further fossil data and exploration of the factors affecting Bayesian tip dating will lead to better resolution of eutherian evolutionary history.

## Conclusions

Inspired by the taxon-sampling strategies that have been adopted by a range of genome-sequencing initiatives, we have investigated the impacts of increasing taxon sampling on Bayesian tip dating under the unresolved FBD process. Our results have shown that denser taxon sampling does not always lead to better posterior estimates of evolutionary parameters in this framework. Instead, performance is also influenced by rate variation among lineages, the positions of fossil taxa, the prior on the root age, and model misspecification. We recommend that future studies should explore additional simulation conditions, such as lower sampling fractions and higher fossil recovery rates. Analysing a wider range of phylogenomic data sets will also provide further insights into the benefits of denser taxon sampling in practice. With a greater understanding of the interplay between taxon sampling and other factors, Bayesian molecular dating can be applied more effectively to phylogenomic data sets to resolve evolutionary timescales.

## Acknowledgements

We acknowledge TianHe-1 (A) at National SuperComputer Center in Tianjin, China for providing computing resources that have contributed to the most results reported in this paper, Chuan Wang for technical supports on computation, and Ming-Qiang Wang for providing scripts on density scatter plots.

